# Large-scale single-virus genomics uncovers hidden diversity of river water viruses and diversified gene profiles

**DOI:** 10.1101/2024.04.18.589877

**Authors:** Yohei Nishikawa, Ryota Wagatsuma, Yuko Tsukada, Lin Chia-ling, Rieka Chijiiwa, Masahito Hosokawa, Haruko Takeyama

**Affiliations:** Computational Bio Big-Data Open Innovation Laboratory, AIST-Waseda University, 3-4-1 Okubo, Tokyo, 169-0082, Japan; Research Organization for Nano & Life Innovation, Waseda University, 513 Waseda tsurumaki-cho, Tokyo, 162–0041, Japan; Department of Life Science and Medical Bioscience, Waseda University, 2-2 Wakamatsu-cho, 162–8480, Tokyo, Japan; Institute for Advanced Research of Biosystem Dynamics, Waseda Research Institute for Science and Engineering, Graduate School of Advanced Science and Engineering, Waseda University, 3-4-1 Okubo, Shinjuku-ku, Tokyo 169-8555, Japan

## Abstract

Environmental viruses (primarily bacteriophages) are widely recognized as playing an important role in ecosystem homeostasis through the infection of host cells. However, the majority of environmental viruses are still unknown as their mosaic structure and frequent mutations in their sequences hinder genome construction in current metagenomics. To enable the large-scale acquisition of environmental viral genomes, we developed a new single-viral genome sequencing platform with microfluidic-generated gel beads. Amplification of individual DNA viral genomes in mass-produced gel beads allows high-throughput genome sequencing compared to conventional single-virus genomics. The sequencing analysis of river water samples yielded 1431 diverse viral single-amplified genomes, while viral metagenomics recovered 100 viral metagenome-assembled genomes at the comparable sequence depth. The 99.5% of viral single-amplified genomes were determined novel at the species level, most of which could not be recovered by a metagenomic assembly. The large-scale acquisition of diverse viral genomes identified protein clusters commonly detected in different viral strains, allowing the gene transfer to be tracked. Moreover, comparative genomics within the same viral species revealed that the profiles of various methyltransferase subtypes were diverse, suggesting an enhanced escape from host bacterial internal defense mechanisms. Our use of gel bead-based single-virus genomics will contribute to exploring the nature of viruses by accelerating the accumulation of draft genomes of environmental DNA viruses.

## Introduction

Environmental viruses (primarily bacteriophages) are the most abundant and diverse biological agents [1]. Through infection of host cells, lytic viruses have a major impact on nutrient and energy cycles [2], while lysogenic viruses contribute to the diversification of the host genomes as one of the major mobile genetic elements [3]. Viruses are abundant in aquatic environments (10^4^–10^8^ particles/mL) [4, 5], and various studies have researched the genome sequencing of aquatic DNA viruses. Currently, the most commonly used method for viral genome sequencing is viral metagenomics, which has been applied to various environmental samples, including soil [6], sea sediment [7], and freshwater [8]. While viral metagenomics has collected diverse viral sequences and viral databases are being expanded, the diversity of the environmental viruses and their functions remain unclear [9]. Current metagenomics has difficulty constructing viral genomes because they are highly mosaic, with frequent genomic mutations and recombination, reducing the efficiency of metagenomic assembly [10]. As genome variation can cause critical changes in viral ecology, including determining host ranges [11], the large-scale acquisition of viral genomes and elucidation of their diversity are essential for understanding their functions.

Single-virus genomics, which uses fluorescence-activated cell sorting (FACS) for viral particle isolation and whole genome amplification (WGA), is an alternative method for obtaining viral sequences, including genomic microdiversity [12]. To date, collecting viral single-amplified genomes (vSAGs) has revealed double-stranded (ds) DNA viral populations dominating oceans, which have been overlooked by metagenomics [13]. However, isolating nanometer-sized viral particles using FACS is technically challenging, and the efficiency of genome amplification was limited to 20% of sorted samples at most [13–15]. Therefore, the number of recovered vSAGs has been limited, and only the most abundant populations were recovered [16].

This study proposes a new single-virus genome sequencing platform using microfluidic-generated gel beads. Microfluidic gel beads have been used in diverse studies as reaction vessels for high-throughput single-cell analysis, including genome amplification of prokaryotic cells [17–20]. Gel beads encapsulate viral particles in a high-throughput manner, allowing genome amplification of > 10^5^ viruses in a single tube at a time. Following WGA, gel beads with amplified DNA were isolated by FACS, which improves the sorting efficiency compared to the conventional FACS sorting of viral particles. Validation using model viruses showed that high-throughput single-virus genomics provides accurate sequence information. In addition, we demonstrate the power of this platform to uncover the genomic diversity of river water viruses.

## Results

### Gel bead-based single-virus genomics enables high-throughput and accurate viral genome sequencing

Viruses were encapsulated into 30 µm-diameter microfluidic gel beads at the single-particle level, followed by capsid lysis and WGA. This platform applies modified protocols of the previously reported bacterial single-cell genome sequencing method (single amplified genomes in gel beads: SAG-gel [21, 22]) for viruses. As proof of principle, viruses packaging two different types of DNA (Lambda and Charomid 9-42) were prepared. Gel bead-based WGA was performed using a mixture of the two viruses (Fig. 1A). After three hours of WGA, DNA amplification was confirmed in gel beads (Fig. 1B). The average fluorescence-positive rate was 3.2%. The viral concentration was calculated to be 9.4 × 10^7^ pfu/µg (*n* = 3), which was slightly lower than the kit reference value (1 × 10^8^ pfu/µg for standard Lambda DNA). In contrast, when we encapsulated the packaging extract without DNA packaging as a control, the fluorescence-positive rate was less than 0.1% (*n* = 3). Next-generation sequencing of amplified DNA in gel beads yielded a median of 186 K trimmed sequence reads for 104 quality-filtered samples, 92.3% (96/104) of which predominantly contained sequence reads mapped to the reference. In the remaining 7.7% (8/104) of samples, 89.4 ± 2.3% of sequence reads were mapped to the *Escherichia coli* genome. Since the sequence results of control samples without DNA packaging also showed a predominance of *E. coli* reads, we consider these sequences to be derived from the packaging extract. All sequenced gel beads had 95.5 ± 5.9% of their quality-filtered sequence reads mapped preferentially to either Lambda DNA or Charomid DNA, with only four gel beads containing > 1% of the other viral sequence reads (Fig. 1C and Table S1). This result suggests that the risk of contamination of the other viral sequences could be reduced by encapsulating dispersed viral particles at appropriate concentrations. The genome coverage was 84.8 ± 24.6% (*n* = 90) for Lambda DNA and 96.6 ± 3.8% (*n* = 6) for Charomid DNA (Fig. 1D). A preliminary estimation of the two viral concentrations showed the number of viral particles containing Lambda DNA was 8.2 - 12 times larger than that of Charomid DNA. The sequencing results showed 90 gel beads were attributed to Lambda DNA and six gel beads were attributed to Charomid DMA, a ratio of 15 times larger. We then employed the Gini coefficient to evaluate the amplification bias [23, 24]. The average Gini coefficient was 0.38 for Lambda DNA and 0.088 for Charomid DNA. These values suggest that our platform showed a more uniform amplification across the viral genomes than bacterial single-cell genome amplification methods with higher Gini coefficients observed [25].

**Fig. 1.**
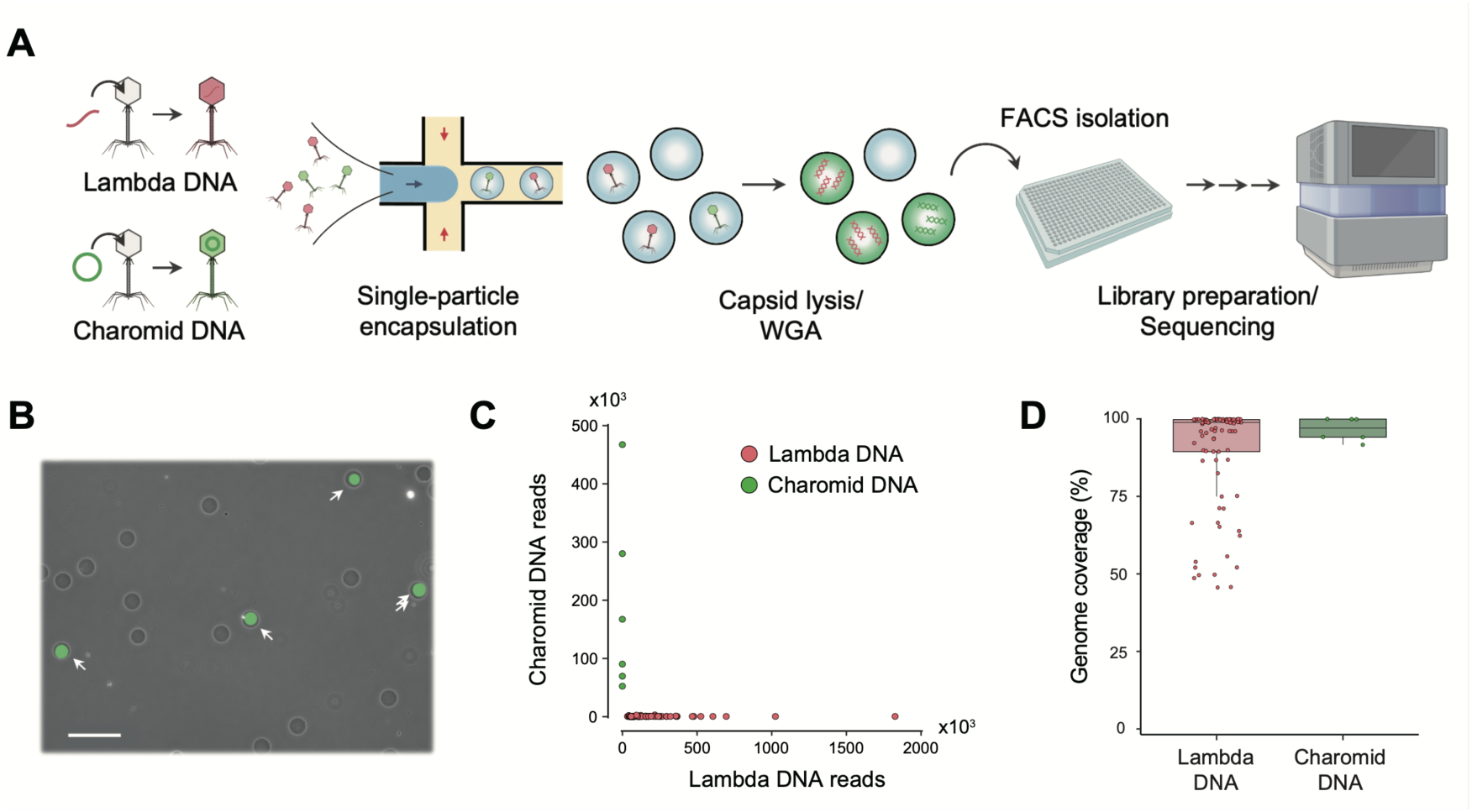
Gel beads enable high-throughput WGA of viruses at the single-particle level. (A) Workflow of single-virus genomics. Viruses are encapsulated into 30-μm of microfluidic droplets with ultra-low melting temperature agarose. After the solidification of agarose, each virus captured in a gel bead proceeded to undergo capsid lysis and WGA. Gel beads with amplified DNA are isolated with FACS and proceeded to the library preparation and next-generation sequencing. (B) Microscopic image of gel beads after WGA. The amplified viral DNA is stained with SYBR Green I, with a scale bar of 100 μm. (C) Comparison of the number of sequence reads mapped to the Lambda and Charomid reference sequences. Samples were labeled with the reference with the most sequence reads to be mapped. Only four of the 96 gel beads contained > 1% of the other viral sequence reads, suggesting a low risk of DNA contamination. (D) Genome coverage by the sequence reads derived from each gel bead

### Large-scale single genome sequencing of viruses from river water

We collected surface river water from an urban river in Tokyo and prepared viral fractions using two methods: (Ⅰ) suction filtration [26] and (Ⅱ) flocculation with FeCl_3_ [27]. These two methods are commonly used for virus particle recovery, and their recovery efficiencies were examined. TEM (transmission electron microscopy) images of the viral fractions showed various morphologies of virus-like particles, including those with tail structures (Fig. 2A).

**Fig. 2.**
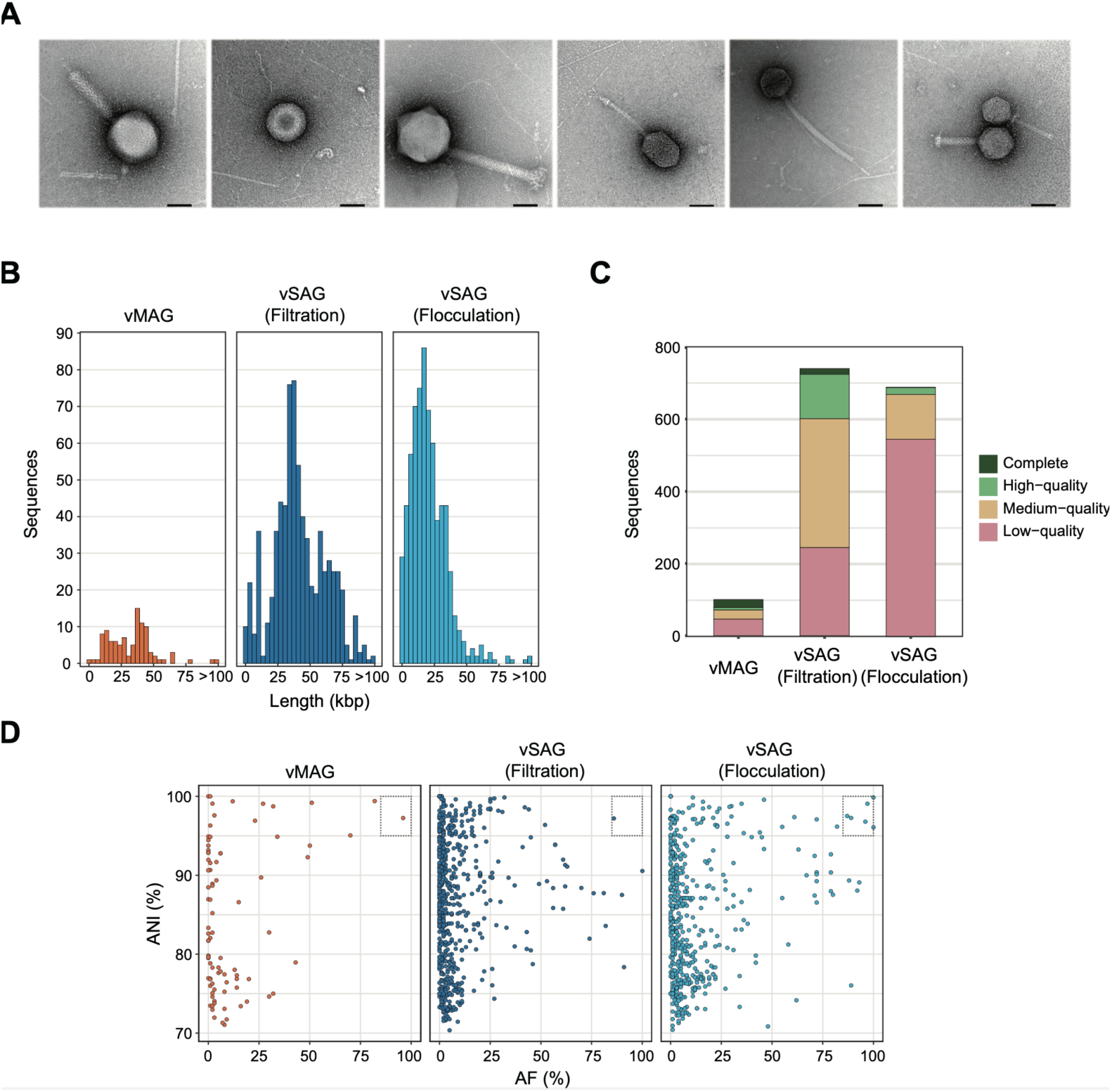
Sequence quality evaluation of vSAGs and vMAGs collected from river water. (A) Negative staining TEM images of viral suspensions were taken at a magnification of 100000×. Various shapes of virus-like particles were recovered from river water. Most particles are dispersed, while some seem to be physically attached. The scale bar is 500 nm. Viruses were collected using two different methods ((Ⅰ) suction filtration and (Ⅱ) flocculation with FeCl_3_), and single-virus genomics was performed with each viral suspension. Metagenomic DNA was extracted from the viral suspensions collected by method (Ⅰ). After next-generation sequencing and the construction of vSAGs and vMAGs, (B) the sequence length, (C) the collected number and the estimated quality by CheckV, and (D) the sequence similarity against reference sequences in the IMG/VR v3 database were evaluated. If the average nucleotide identity (ANI) was > 95% and the alignment fraction (AF) was > 85%, the viral sequences were judged to be known sequences (dashed square).

Following WGA, the fluorescence-positive rate of gel beads was ≤ 13.2% and the calculated concentration of DNA virus in the river water was 9.2 × 10^5^ particles/mL by method (Ⅰ) and 2.6 × 10^5^ particles/mL by method (Ⅱ). A total of 1536 DNA-amplified gel beads (768 for each method) were isolated and proceeded to next-generation sequencing, yielding 12.4 Gb sequence reads by method (Ⅰ) and 11.6 Gb by method (Ⅱ), respectively. While genome sequencing of environmental viruses poses a risk of contamination, such as physical aggregation of viral particles, there are currently no analytical tools to assess contamination in viral sequences. Therefore, we employed the metagenomic binning tool to construct vSAGs to exclude potentially contaminating sequences. If contigs derived from a single gel bead were classified into multiple distinct bins, contigs in the bin with the highest completeness were used to construct the vSAG, and contigs in the other bins were excluded. We found 54.2% (775/1431) of the gel beads had contigs classified into multiple distinct bins. It should be noted that excluded contigs do not directly indicate viral sequence contamination since contigs derived from one viral species could be classified into multiple bins due to the lack of sequence information. Finally, we recovered 1431 vSAGs from 740 samples (96.3% of the total) by method (Ⅰ) and 691 samples (90.0% of the total) by method (Ⅱ), demonstrating a significantly higher recovery rate than conventional single-virus genomics [13–15]. Of the 105 samples with no viral sequences detected, 54 had small data sizes (less than 5,000 paired reads), and most of the remaining sequences were determined to contain fragmented reads of bacteria or viruses. In the metagenomic sequencing that uses extracted DNA from the viral suspension in method (Ⅰ), we recovered 16.8 Gb reads and 100 viral metagenome-assembled genomes (vMAGs). It is noted that we used different processes for constructing vSAGs and vMAGs.

The number of vSAGs or vMAGs per 1 Gb of sequence reads for method (Ⅰ), method (Ⅱ), and for metagenomics was 59.7, 59.6, and 5.95, respectively. The median length of vSAGs was 39781 bp for method (Ⅰ) and 18296 bp for method (Ⅱ), and that of vMAGs was 34985 bp (Fig. 2B and Table S2). In the quality assessment by CheckV [28], 496 (67.0 %) of vSAGs by method (Ⅰ), 144 (20.8%) of vSAGs by method (Ⅱ), and 54 (54.0 %) of vMAGs were medium- or high-quality (Fig. 2C and Table S2). No viral sequence was detected in 6.8% (105/1536) of gel beads.

As a result of referring to IMG/VR v3 database, 99.9% (739/740) and 99.1% (685/691) of vSAGs with method (Ⅰ) and method (Ⅱ), respectively, were determined novel at the species level, with only seven sequences corresponding to the reference (Fig. 2D). In the vMAGs, 99.0% (99/100) were determined novel. Of the two virus concentration methods, method (Ⅰ) exhibited a longer median length and higher genome quality than method (Ⅱ), consistent with a previous study [27].

### Single-virus genomics recovers highly diverse and low-abundance viral sequences

Phylogenetic diversity of vSAGs and vMAGs was evaluated by constructing viral clusters (VCs) based on a protein-sharing network (Fig. 3A and S1), consistent with the genus-subfamily level classification. In total, we detected 289 VCs from 873 vSAGs (223 VCs from method (Ⅰ) and 153 VCs from method (Ⅱ)) and 38 VCs from 50 vMAGs (Table S2). There were 27 VCs detected in common between vSAGs and vMAGs. The number of VCs clustered with the reference sequence was only 14 in vSAGs and seven in vMAGs, suggesting that numerous novel sequences at the genus-subfamily level were obtained.

**Fig. 3.**
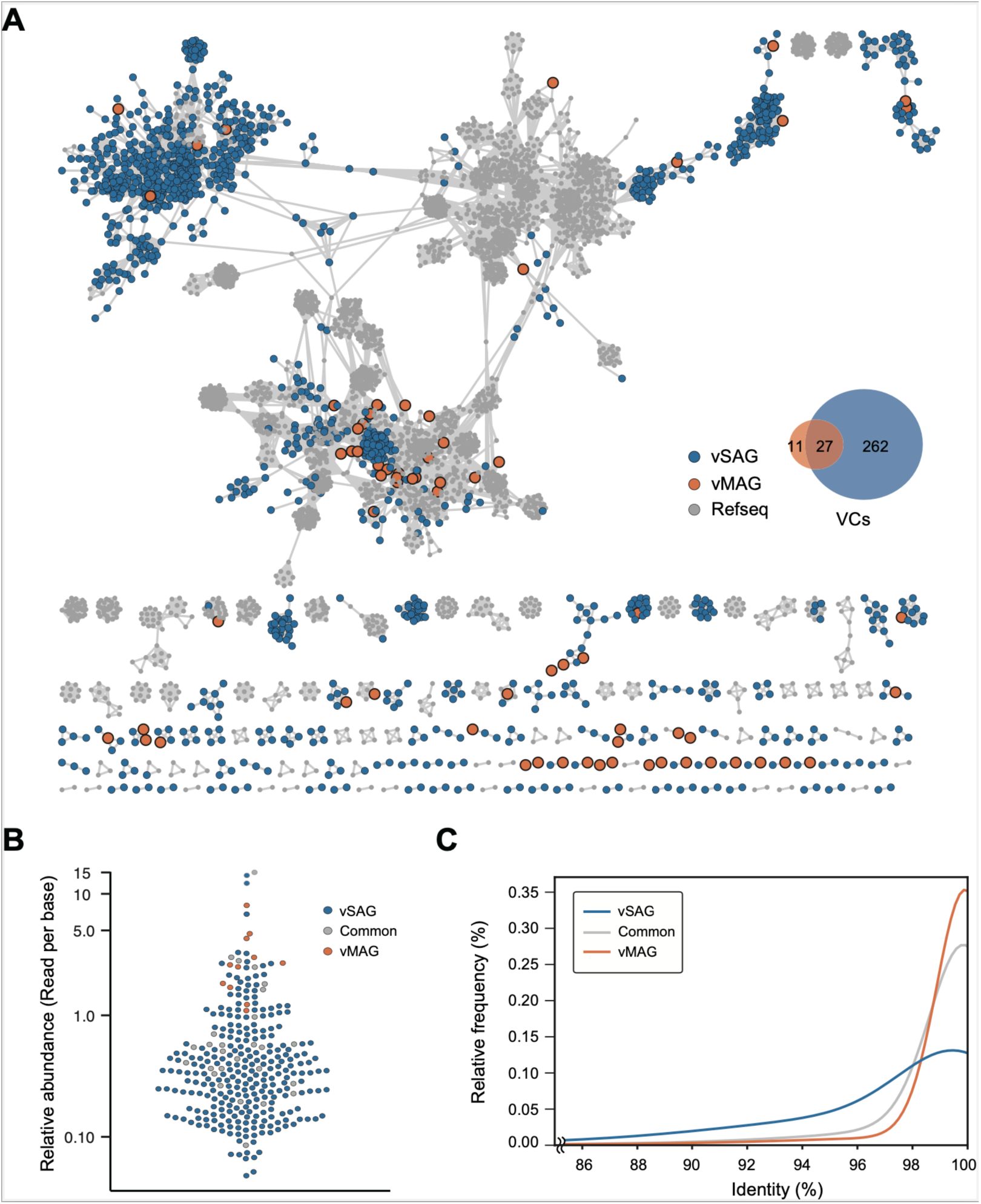
Protein-sharing network of 1431 vSAGs and 100 vMAGs. (A) Protein-sharing network analysis of 1431 vSAGs and 100 vMAGs with RefSeq reference database using vConTACT2. Blue dots represent vSAGs, and orange dots represent vMAGs. Gray dots are references. Only the sequences sharing at least one protein cluster with other sequences are represented. (B) Relative abundance of vSAGs and vMAGs. The vertical axis is the number of mapped metagenomic sequence reads to the representative sequences, expressed as the logarithm of reads per base. (C) Metagenomic sequence reads recruitment patterns (also referred to as diversity curves) for the representative sequences of each VC. Curves represent the percentage of recruited sequence reads at each nucleotide identity value [13]

Taxonomic classification was conducted using the methodology employed in the IMG/VR v3 database, which references the 2019 ICTV Release. Of the 50 vMAGs that clustered into VCs, 20% (10/50) were taxonomically assigned at the family level, all of which were tailed dsDNA viruses (*Caudoviricetes*), including *Siphoviridae*, *Myoviridae*, and *Podoviridae* (Fig. S2A and Table S2). In contrast, only 6.5% (57/873) of vSAGs clustered into VCs were taxonomically assigned. Most sequences were assigned to dsDNA viruses (Fig. S2B), whereas two vSAGs were assigned as *Mimiviridae* and four as *Lavidaviridae,* which are giant viruses and virophages, respectively. In addition, ten single-stranded (ss) DNA viruses, including *Circoviridae* and *Microviridae*, were recovered because ssDNA was also amplified by WGA [29]. Of the 10737 genes hit by the RefSeq viral protein, 10324 (96.2%) were best-matched to bacteriophage genes, suggesting that most of the viral sequences were from bacteriophages.

We then assessed the relative abundance of vSAGs and vMAGs in the metagenomic raw reads. vSAGs contained viral sequences of diverse abundance, including relatively low abundance viral sequences (Fig. 3B). Species-specific recruitment patterns for each VC [13] indicated that vSAGs recovered viral populations with both microdiversity and macrodiversity while vMAGs had a relatively low sequence diversity (Fig. 3C). Since the metagenomic raw reads contained 68.9% (986/1431) of the vSAG sequences with > 50% coverage, we suggest that most of the viral sequences were missed during the process of metagenomic assembly.

### Identification of auxiliary metabolic genes shared among viruses of different lineages

We detected viral auxiliary metabolic genes (AMGs) from vSAGs and vMAGs that were of medium- or high-quality and clustered into VCs to assess how viral infection modulates host metabolism. We detected 280 AMGs of 70 types in 161 vSAGs and 16 AMGs of 10 types in 11 vMAGs (Table S3). The highest number of AMGs per vSAG was 10, and the most abundant functional category of AMGs was affiliated with organic nitrogen metabolism (Fig. S3A, B).

The protein cluster (PC) profiles were then evaluated to determine gene transfer among viruses. We detected 9754 PCs from 494 vSAGs and 34 vMAGs with medium- or high-quality. Although a previous report using metagenomic contigs showed that distinct VCs share few protein groups [30], large-scale single-virus genomics revealed that 30.7% (2993/9754) of PCs were shared in multiple distinct VCs (Table S4). There were 233 AMGs of 33 types from multiple VCs, which corresponds to 2.27% (68/2993) of PCs (Fig. 4A). In contrast, 47 AMGs of 37 types were detected from only a single VC, which corresponds to 0.84% (57/6761) of PCs. These results suggest that AMG-containing PCs were more frequently distributed among VCs (*p* = 0.01, Fisher’s exact test). Next, a phylogenetic analysis of vSAGs harboring BTLCP, the most frequently identified AMG, was conducted. The proteomic tree created by ViPTree showed clades corresponding to each VC (Fig. 4B). In addition, the phylogenetic tree of the terminase large subunit (TerL), which is often used in phylogenetic analysis, showed clades corresponding to each VC with robust bootstrap support (Fig. 4C). In contrast, the phylogenetic tree of BTLCP genes did not cluster by VCs but branched within diverse clades (Fig. 4D). As some clades were supported by sufficient bootstrap values (SH-like aLRT ≥ 80%), BTLCP may have been acquired via horizontal gene transfer after the divergence of these VCs.

**Fig. 4.**
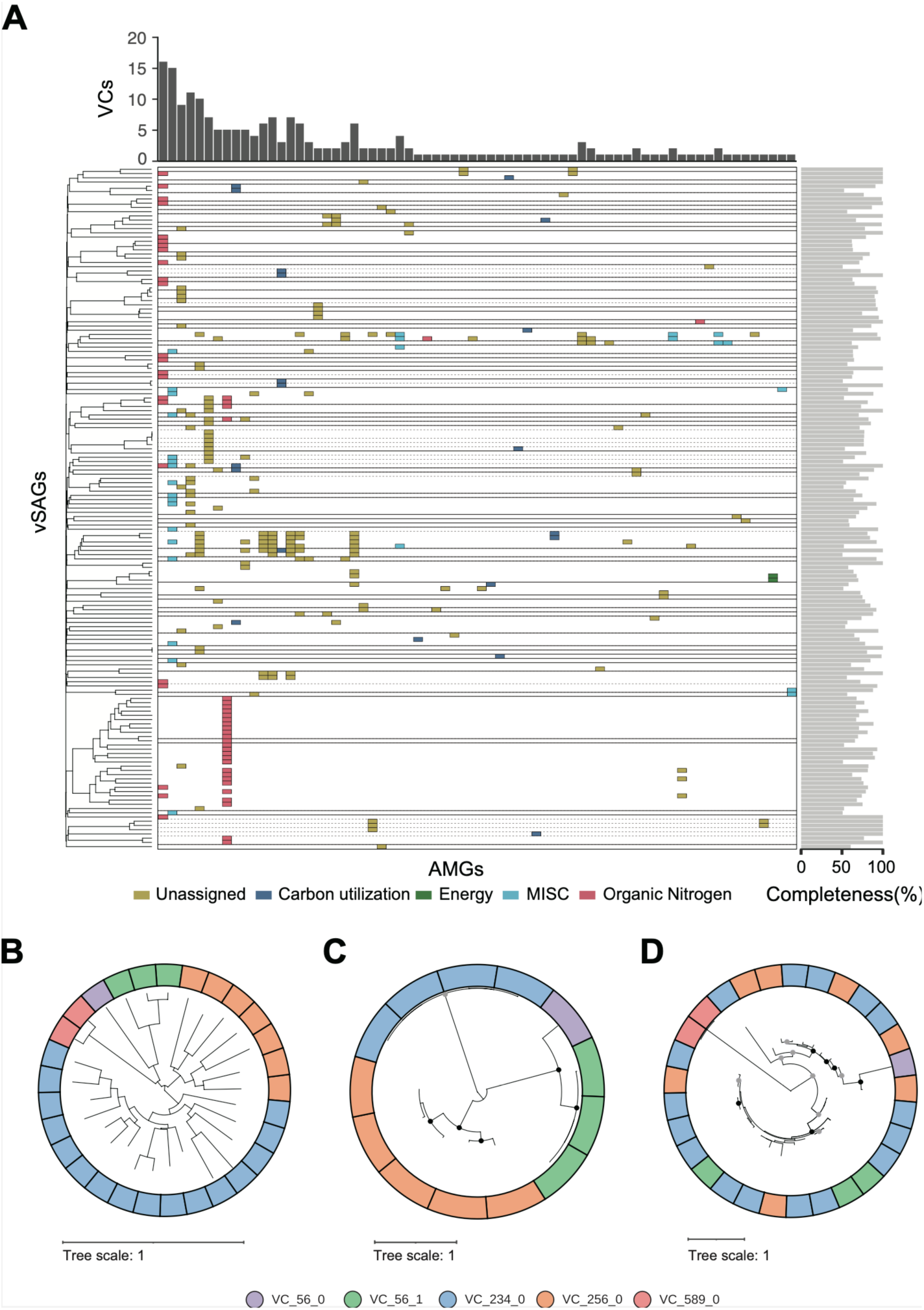
Identification of AMGs commonly detected from multiple VCs. (A) The distribution profiles of AMGs among vSAGs. Each row represents a single vSAG, and each column represents a single AMG. Straight lines separate each VC, and subclusters within a VC are separated by dashed lines. The color of the panel distinguishes the AMG categories determined by DRAM-v. The left side of the figure shows the phylogenetic tree of vSAGs, and the right-side bar plot shows the completeness of each vSAG. The histogram at the top shows the number of VCs containing each AMG, showing 33 types of AMGs from multiple VCs and 37 types of AMGs from a single VC. (B) Proteomic tree created by ViPTree of vSAGs with BTLCP detected. Maximum-likelihood phylogenies of (C) TerL and (D) BTLCP. Branches with bootstrap support by SH-like aLRT higher than 80% and ultrafast bootstrap 95% are indicated by black dots. If those branches were only supported by SH-like aLRT higher than 80%, they are shown in grey. Each color of the outer circle represents a different VC value.

### Comparative genomics reveals the profiles of methyltransferase (MTase) are diversified within the same viral species

Clustering was performed at the vOTU (viral operational taxonomic unit) level, which corresponds to the viral species level, to assess the viral genomic microdiversity. Based on the Minimum Information about an Uncultivated Virus Genome (MIUViG) criteria [31], 1197 vOTUs were generated. Of these, 11.5% (138/1197) of vOTUs consisted of multiple viral sequences. We focused on vOTU572, obtained in one from vMAG with a completeness of 51.0% and six from vSAGs with an average completeness of 93.7%. vOTU572 is novel at the genus level compared with the IMG/VR v3 database [32] and forms an independent VC. The seven sequences within vOTU572 were very similar, as the sequence identity of the TerL was 100%, and ANI was > 98.92%. In contrast, the profile of orthologous groups revealed the presence of two genomic regions: highly conserved regions (core) and regions with structural diversity (flexible) (Fig. 5A). Most of the regions recovered by the vMAG belonged to the core regions, and the flexible regions were lost. While metagenomic raw reads were also mapped to flexible regions (Fig. S6), the most flexible region was not detected in metagenomic contigs. In addition, even when only metagenomic reads corresponding to the flexible regions were extracted and re-assembled, the flexible region was not constructed. These results suggest that the reads corresponding to the flexible region were sequenced but missing during assembly. Genes were annotated in 11.9% (5/42) of the core regions and 22.6% (7/31) of the flexible regions. The core region included genes with TerL, bacteriophage lambda NinG protein (PF05766), and putative peptidoglycan binding domain (PF01471), while we identified multiple methyltransferase (MTase) subtypes in the flexible regions. The sequence alignment of each vSAG revealed that various MTase subtypes were inserted into the genomes (Fig. 5B). As the viral MTase contributes to escape from the bacterial restriction-modification systems by methylating their DNA [33], diversifying the pattern of MTase by frequent homologous recombination suggests a strain-level adaptation strategy of viruses against the internal immune mechanisms of host bacteria.

**Fig. 5.**
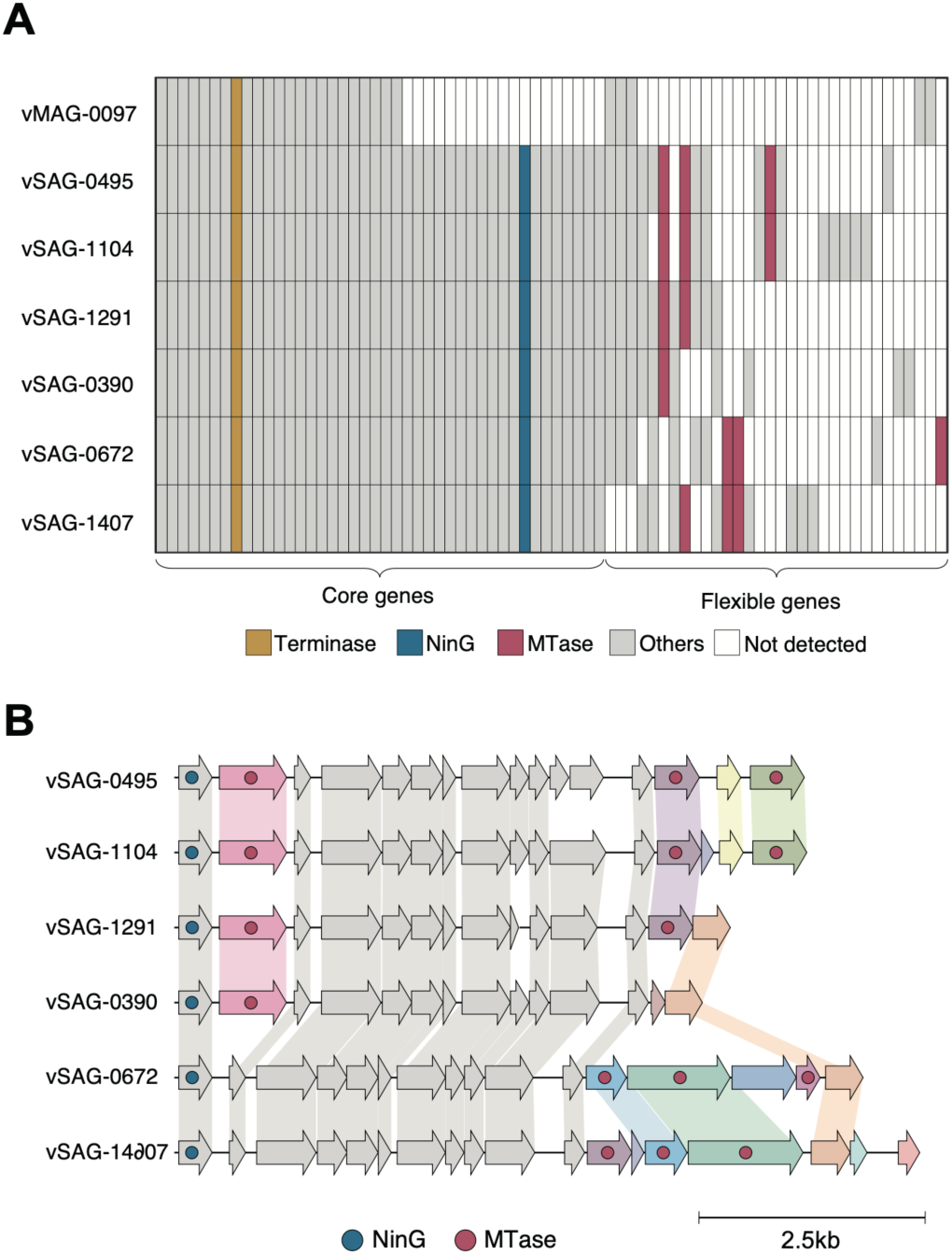
Comparative genomics within the same vOTU (vOTU572) (A) An orthologous groups matrix where each row represents a single orthologous group and columns represent six vSAGs and one vMAG grouped into vOTU572. The top sequence is derived from the vMAG, and the others are derived from vSAGs. The presence or absence of a gene is indicated by color. Gray columns indicate genes that were not annotated, and white columns indicate the genes that were not detected. It is noted that we used different processes for constructing vSAGs and vMAGs. (B) Alignment of six vSAGs partial genomes. Open reading frames are represented by block arrows. Gray-colored arrows indicate genes in the core region, while other colored arrows indicate genes in the flexible region. The different colors of the genes in the flexible region indicate different orthologous groups, showing that six different MTase subtypes were detected in vOTU572. The lead depth in the flexible region of vSAGs was 258.8 on average and 23.3 at the lowest.

## Discussion

Environmental viruses are more diverse than environmental bacteria, and genomic mutations occur frequently, leaving many challenges for high-resolution genome analysis [9]. Single-virus genome sequencing using FACS has been performed; however, it has some difficulties in sorting accuracy, and previous reports have recovered only a few dozen viral sequences mainly because of the low success rate of WGA [13–15]. Here, we developed a gel bead-based single-virus genomics platform and recovered a total of 1431 viral sequences from 93.2% of the isolated gel beads. Our platform dramatically improved the efficiency and throughput of viral genome sequencing and recovered a large number of diverse viral sequences, including those with low relative abundance. On the other hand, the results of mapping metagenomic raw reads to vSAGs imply that metagenomic assembly is hampered in vMAG construction by viral genomic microdiversity, which is recognized as “the great metagenomics anomaly” [10, 34]. As 91.8% of the viral sequences obtained in this study were novel at the family level, accumulating vSAGs could help in the phylogenetic classification of environmental viruses.

High-throughput genome sequencing enabled high-resolution detection of PCs commonly detected in different VCs, allowing the tracing of gene distribution among viruses. As a higher percentage of AMGs were commonly detected in multiple distinct VCs compared to other PCs, these AMGs are suggested to play a significant role in virus survival. For example, sequence homology analysis suggested that BTLCP can be exchanged via co-infection with the same bacterial host and transferred among distinct VCs. As BTLCP is suggested to be involved in catalyzing post-translational protein modification [35], modifying prokaryotic surface structures [35], and biofilm formation [36], the acquisition of such AMGs could be advantageous for host bacterial adaptation to the environment.

Large-scale single-virus genomics has enabled obtaining closely related but diverse viral sequences. Comparative genomics revealed frequent homologous recombination of various MTase subtypes within the same vOTU. A recent study reported that viral host specificity is not determined by proteins involved in viral adhesion to the cell membrane but rather by the internal defense mechanisms of the host bacteria [37]. Furthermore, a study analyzing nearly identical viral genomes from distant aquatic ecosystems also reported frequent transfer of MTase and suggested that MTase protects viruses against host-encoded restriction-modification systems [38]. Our results imply that viruses with multiple MTase subtypes could enhance their ability to escape bacterial internal defense mechanisms, broadening their host ranges. It has been reported that genes involved in viral immunity are frequently horizontally transferred among bacteria [39]. Correspondingly, this study confirmed that genes suggested to be involved in host adaptation are widely distributed among viruses.

We developed a gel bead-based single-virus genomics platform to reveal gene transfer among diverse viruses and microdiversity within the same vOTU in river water viruses. As our platform can recover various single-virus genome sequences, large-scale comparative genomics has enabled the detection of genetic recombination, which is the most significant factor in virus diversification [38, 40]. In contrast, 39% (558/1431) of the acquired vSAGs did not cluster into a VC, suggesting that larger genome sequencing would be required for a comprehensive protein-sharing network analysis. Although the current platform allows obtaining > 1400 viral sequences in a single sequence run, further development, such as multiplexed barcoding [41], will be necessary. Also, whole genome amplification using phi29 may experience amplification bias for genomes with varying sizes/linearity/GC content. In addition, since the contigs obtained from a single gel bead are usually fragmented, the protocol for constructing vSAGs needs to be improved. Bioinformatics tools specialized for single virus genomics should be developed to evaluate the viral contamination. In this study, we performed a snapshot analysis of the viral community; however, time-lapse sampling would allow the analysis of the temporal variation of the viral community and horizontal gene transfer occurring at that moment. Although viral genome sequencing with long-read sequencers has been demonstrated, it requires a substantial amount of water to extract DNA of sufficient quality [42, 43]. In contrast, our platform can perform genome sequencing from a few hundred milliliters of water, making it easier to perform analyses on temporal variation in viral communities. Furthermore, cross-referencing single-cell genomes of environmental bacteria with vSAGs will enhance our understanding of virus-host interactions, including host range and bacterial immunity against viruses. Our single-virus genomics platform will contribute to elucidating the unknown functions of environmental viruses.

## Acknowledgment

This work was supported by MEXT KAKENHI grant number 17H06158, 21H01733, and JST ACT-X Grant Number JPMJAX20BE. The Human Genome Center (University of Tokyo) provided supercomputing resources. TEM imaging was performed at Hanaichi Ultra Structure Research Institute. Part of Fig. 1A was created using BioRender.com.

---

**Fig. S1.**
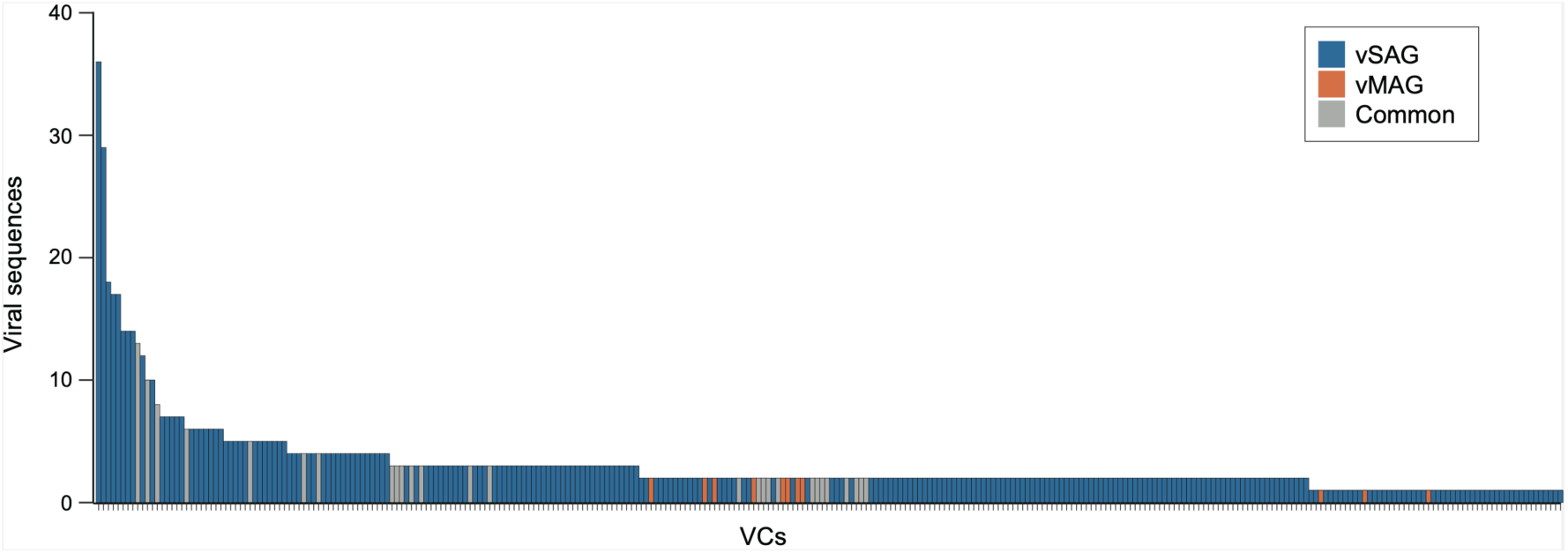
The number of viral sequences distributed in each VC

**Fig. S2.**
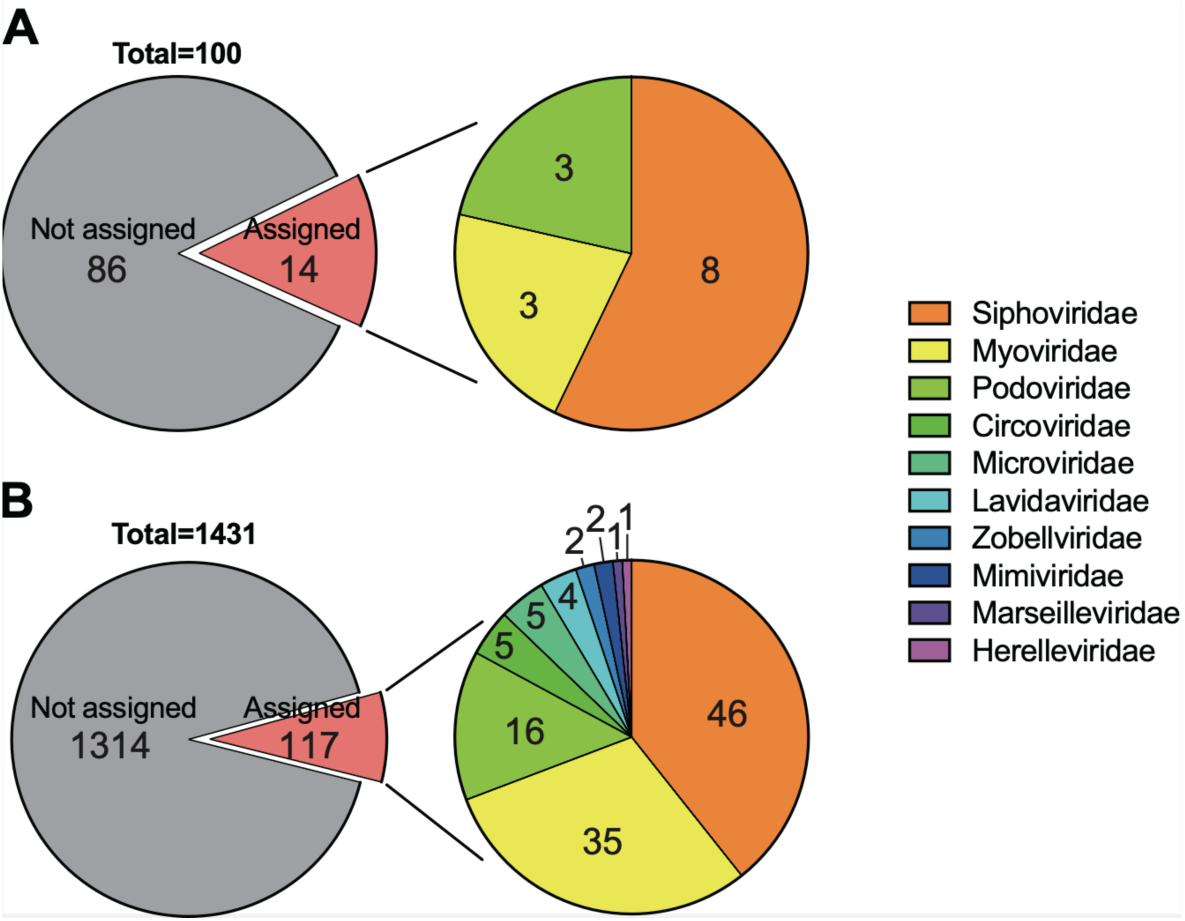
Taxonomic assignment of (A) 100 vMAGs and (B) 1431 vSAGs. Taxonomic classification was conducted using the methodology employed in the IMG/VR v3 database, which references the 2019 ICTV Release.

**Fig. S3.**
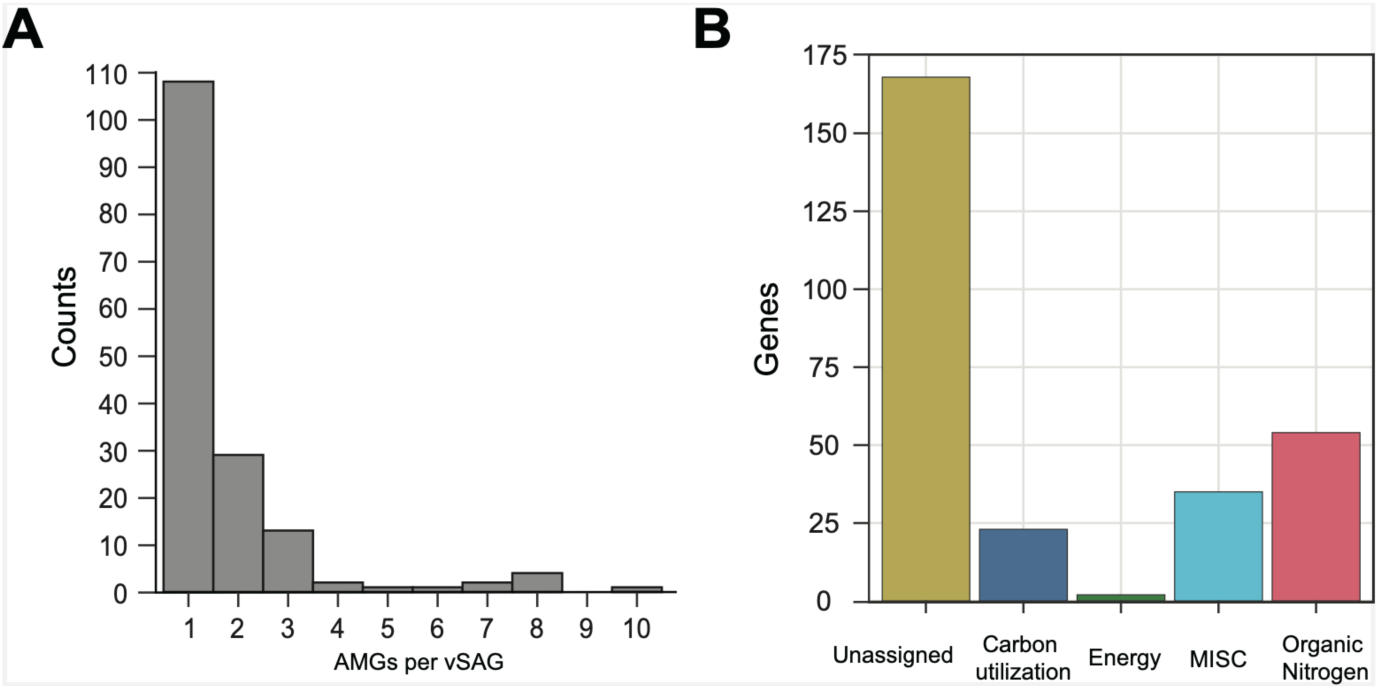
Detection of AMGs from vSAGs. (A) Histogram of the number of AMGs for each vSAG (B) The number of detected AMGs for each category defined by DRAM-v

**Fig. S4.**
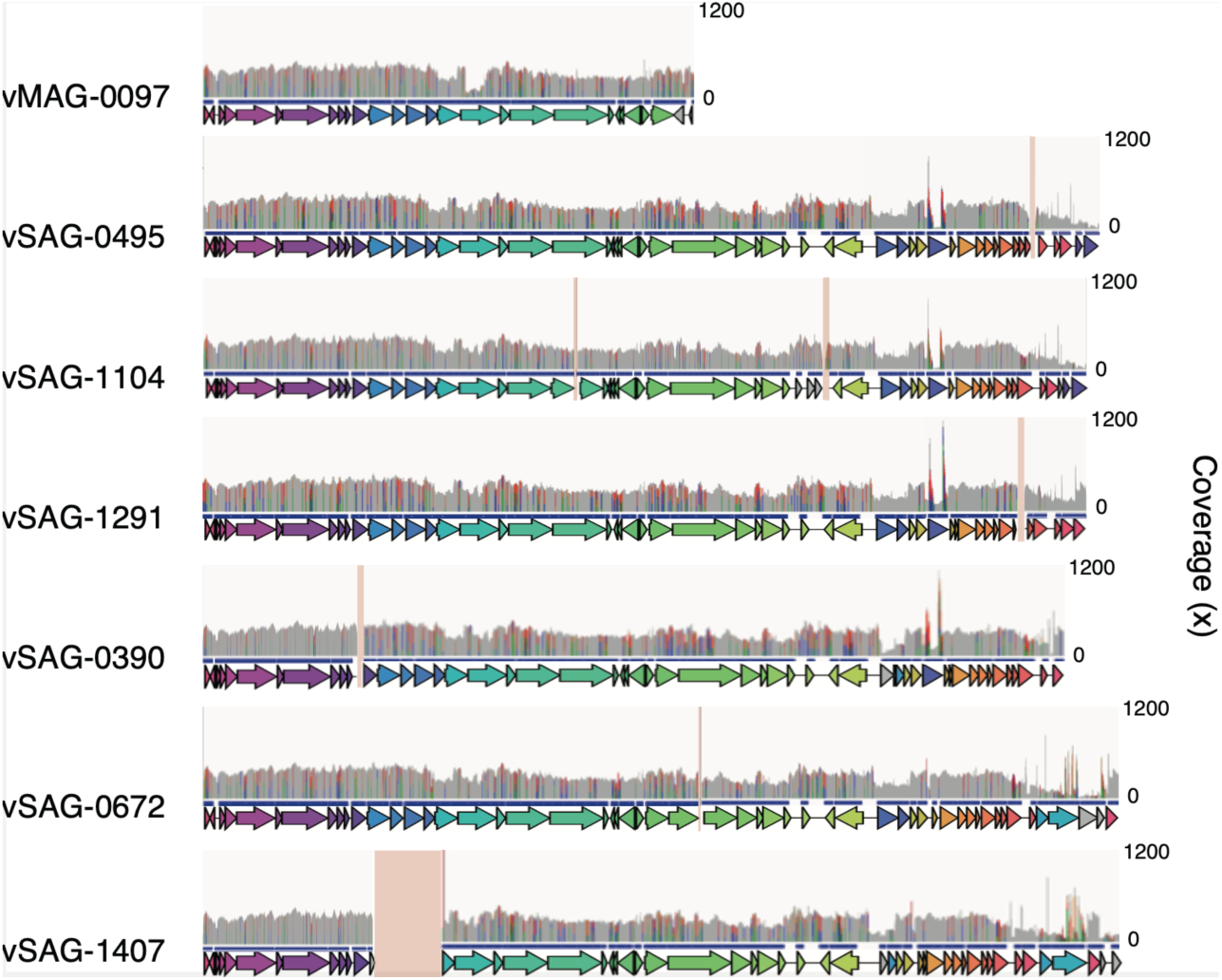
Metagenome read mapping to one vMAG and six vSAGs of vOTU572. Metagenome sequence reads were mapped to one vMAG and six vSAGs using bwa-mem. The y-axis shows the read depth; the upper limit was 1200. The gap region is indicated by the pink bars. As metagenomic reads were also mapped in regions of vSAGs that were not constructed in the vMAG, it suggests that sequence information was missing during metagenome assembly.

**Table S1 The number of sequence reads mapped to the Lambda and Charomid reference sequences**

**Table S2 Sequence statistics of vSAGs and vMAGs Table S3 Summary of AMGs**

**Table S4 Summary of PCs detected from each VC**

A table shows the presence patterns of PCs in vSAGs as identified by vConTACT2. The table lists vSAGs with one or more PCs classified as medium- or high-quality.

## References

[1] Rohwer F, Prangishvili D, Lindell D (2009). Roles of viruses in the environment. Environ Microbiol 11: 2771–2774.

[2] Suttle CA (2007). Marine viruses--major players in the global ecosystem. Nat Rev Microbiol 5: 801–812.

[3] Canchaya C, Fournous G, Chibani-Chennoufi S, Dillmann M-L, Brüssow H (2003). Phage as agents of lateral gene transfer. Current Opinion in Microbiology 6: 417–424.

[4] Breitbart M, Bonnain C, Malki K, Sawaya NA (2018). Phage puppet masters of the marine microbial realm. Nat Microbiol 3: 754–766.

[5] Wommack KE, Colwell RR (2000). Virioplankton: viruses in aquatic ecosystems. Microbiol Mol Biol Rev 64: 69–114.

[6] Emerson JB, Roux S, Brum JR, Bolduc B, Woodcroft BJ, Jang HB et al (2018). Host-linked soil viral ecology along a permafrost thaw gradient. Nat Microbiol 3: 870–880.

[7] Rambo IM, Langwig MV, Leao P, De Anda V, Baker BJ (2022). Genomes of six viruses that infect Asgard archaea from deep-sea sediments. Nat Microbiol 7: 953–961.

[8] Moon K, Jeon JH, Kang I, Park KS, Lee K, Cha CJ et al (2020). Freshwater viral metagenome reveals novel and functional phage-borne antibiotic resistance genes. Microbiome 8: 75.

[9] Roux S, Hallam SJ, Woyke T, Sullivan MB (2015). Viral dark matter and virus-host interactions resolved from publicly available microbial genomes. Elife 4.

[10] Ramos-Barbero MD, Martin-Cuadrado AB, Viver T, Santos F, Martinez-Garcia M, Anton J (2019). Recovering microbial genomes from metagenomes in hypersaline environments: The Good, the Bad and the Ugly. Syst Appl Microbiol 42: 30–40.

[11] Paterson S, Vogwill T, Buckling A, Benmayor R, Spiers AJ, Thomson NR et al (2010). Antagonistic coevolution accelerates molecular evolution. Nature 464: 275–278.

[12] Martinez Martinez J, Martinez-Hernandez F, Martinez-Garcia M (2020). Single-virus genomics and beyond. Nat Rev Microbiol 18: 705–716.

[13] Martinez-Hernandez F, Fornas O, Lluesma Gomez M, Bolduc B, de la Cruz Pena MJ, Martinez JM et al (2017). Single-virus genomics reveals hidden cosmopolitan and abundant viruses. Nat Commun 8: 15892.

[14] Martinez-Hernandez F, Fornas O, Martinez-Garcia M (2022). Into the Dark: Exploring the Deep Ocean with Single-Virus Genomics. Viruses 14.

[15] Allen LZ, Ishoey T, Novotny MA, McLean JS, Lasken RS, Williamson SJ (2011). Single virus genomics: a new tool for virus discovery. PLoS One 6: e17722.

[16] Gregory AC, Zayed AA, Conceicao-Neto N, Temperton B, Bolduc B, Alberti A et al (2019). Marine DNA Viral Macro- and Microdiversity from Pole to Pole. Cell 177: 1109–1123 e1114.

[17] Fitzsimons MS, Novotny M, Lo CC, Dichosa AE, Yee-Greenbaum JL, Snook JP et al (2013). Nearly finished genomes produced using gel microdroplet culturing reveal substantial intraspecies genomic diversity within the human microbiome. Genome Res 23: 878–888.

[18] Lan F, Demaree B, Ahmed N, Abate AR (2017). Single-cell genome sequencing at ultra-high-throughput with microfluidic droplet barcoding. Nat Biotechnol 35: 640–646.

[19] Leonaviciene G, Leonavicius K, Meskys R, Mazutis L (2020). Multi-step processing of single cells using semi-permeable capsules. Lab Chip 20: 4052–4062.

[20] Zheng W, Zhao S, Yin Y, Zhang H, Needham DM, Evans ED et al (2022). High-throughput, single-microbe genomics with strain resolution, applied to a human gut microbiome. Science 376: eabm1483.

[21] Chijiiwa R, Hosokawa M, Kogawa M, Nishikawa Y, Ide K, Sakanashi C et al (2020). Single-cell genomics of uncultured bacteria reveals dietary fiber responders in the mouse gut microbiota. Microbiome 8: 5.

[22] Nishikawa Y, Kogawa M, Hosokawa M, Wagatsuma R, Mineta K, Takahashi K et al (2022). Validation of the application of gel beads-based single-cell genome sequencing platform to soil and seawater. ISME Communications 2.

[23] Hammond M, Homa F, Andersson-Svahn H, Ettema TJ, Joensson HN (2016). Picodroplet partitioned whole genome amplification of low biomass samples preserves genomic diversity for metagenomic analysis. Microbiome 4: 52.

[24] Gonzalez-Pena V, Natarajan S, Xia Y, Klein D, Carter R, Pang Y et al (2021). Accurate genomic variant detection in single cells with primary template-directed amplification. Proc Natl Acad Sci U S A 118.

[25] Hosokawa M, Nishikawa Y, Kogawa M, Takeyama H (2017). Massively parallel whole genome amplification for single-cell sequencing using droplet microfluidics. Scientific Reports 7.

[26] Sun G, Xiao J, Wang H, Gong C, Pan Y, Yan S et al (2014). Efficient purification and concentration of viruses from a large body of high turbidity seawater. MethodsX 1: 197–206.

[27] Langenfeld K, Chin K, Roy A, Wigginton K, Duhaime MB (2021). Comparison of ultrafiltration and iron chloride flocculation in the preparation of aquatic viromes from contrasting sample types. PeerJ 9: e11111.

[28] Nayfach S, Camargo AP, Schulz F, Eloe-Fadrosh E, Roux S, Kyrpides NC (2021). CheckV assesses the quality and completeness of metagenome-assembled viral genomes. Nat Biotechnol 39: 578–585.

[29] Kim KH, Chang HW, Nam YD, Roh SW, Kim MS, Sung Y et al (2008). Amplification of uncultured single-stranded DNA viruses from rice paddy soil. Appl Environ Microbiol 74: 5975–5985.

[30] Kieft K, Zhou Z, Anderson RE, Buchan A, Campbell BJ, Hallam SJ et al (2021). Ecology of inorganic sulfur auxiliary metabolism in widespread bacteriophages. Nat Commun 12: 3503.

[31] Roux S, Adriaenssens EM, Dutilh BE, Koonin EV, Kropinski AM, Krupovic M et al (2019). Minimum Information about an Uncultivated Virus Genome (MIUViG). Nat Biotechnol 37: 29–37.

[32] Roux S, Paez-Espino D, Chen IA, Palaniappan K, Ratner A, Chu K et al (2021). IMG/VR v3: an integrated ecological and evolutionary framework for interrogating genomes of uncultivated viruses. Nucleic Acids Res 49: D764–D775.

[33] Bravo JPK, Aparicio-Maldonado C, Nobrega FL, Brouns SJJ, Taylor DW (2022). Structural basis for broad anti-phage immunity by DISARM. Nat Commun 13: 2987.

[34] Okazaki Y, Nakano S, Toyoda A, Tamaki H (2022). Long-Read-Resolved, Ecosystem-Wide Exploration of Nucleotide and Structural Microdiversity of Lake Bacterioplankton Genomes. Msystems 7.

[35] Ginalski K, Kinch L, Rychlewski L, Grishin NV (2004). BTLCP proteins: a novel family of bacterial transglutaminase-like cysteine proteinases. Trends Biochem Sci 29: 392–395.

[36] Rybtke M, Berthelsen J, Yang L, Hoiby N, Givskov M, Tolker-Nielsen T (2015). The LapG protein plays a role in Pseudomonas aeruginosa biofilm formation by controlling the presence of the CdrA adhesin on the cell surface. Microbiologyopen 4: 917–930.

[37] Hussain FA, Dubert J, Elsherbini J, Murphy M, VanInsberghe D, Arevalo P et al (2021). Rapid evolutionary turnover of mobile genetic elements drives bacterial resistance to phages. Science 374: 488–492.

[38] Bellas CM, Schroeder DC, Edwards A, Barker G, Anesio AM (2020). Flexible genes establish widespread bacteriophage pan-genomes in cryoconite hole ecosystems. Nat Commun 11: 4403.

[39] Bernheim A, Sorek R (2020). The pan-immune system of bacteria: antiviral defence as a community resource. Nat Rev Microbiol 18: 113–119.

[40] Kupczok A, Neve H, Huang KD, Hoeppner MP, Heller KJ, Franz C et al (2018). Rates of Mutation and Recombination in Siphoviridae Phage Genome Evolution over Three Decades. Mol Biol Evol 35: 1147–1159.

[41] Kuchina A, Brettner LM, Paleologu L, Roco CM, Rosenberg AB, Carignano A et al (2021). Microbial single-cell RNA sequencing by split-pool barcoding. Science 371.

[42] Zablocki O, Michelsen M, Burris M, Solonenko N, Warwick-Dugdale J, Ghosh R et al (2021). VirION2: a short- and long-read sequencing and informatics workflow to study the genomic diversity of viruses in nature. PeerJ 9: e11088.

[43] Tedersoo L, Albertsen M, Anslan S, Callahan B (2021). Perspectives and Benefits of High-Throughput Long-Read Sequencing in Microbial Ecology. Appl Environ Microbiol 87: e0062621.

